# Effects of parity and early pregnancy on peripheral blood leukocytes in dairy cattle

**DOI:** 10.1101/2024.05.06.592827

**Authors:** M. I. da Silva, N. Oli, F. Gambonini, T. Ott

## Abstract

Subfertility remains a major problem in the dairy industry. Only 35-40% of high-yielding dairy cows and 55-65% of nonlactating heifers become pregnant after their first service. The immune system plays a critical role in the establishment of pregnancy. However, it can also create challenges for embryo survival and contribute to reduced fertility. We conducted 2 separate experiments to characterize changes in subsets of peripheral blood leukocytes (PBL) and their phenotype over the estrous cycle and early pregnancy in heifers and cows. We used flow cytometry and RT-qPCR to assess protein and mRNA expression of molecules important for immune function. We observed that monocytes and T cells were most affected by pregnancy status in heifers, whereas, CD8^+^ lymphocytes and natural killer (NK) cells were most affected during early pregnancy in cows. Changes in immune parameters measured appeared to be greater in heifers than cows including changes in expression of numerous immune function molecules. To test the hypothesis, we conducted a third experiment to simultaneously analyze the immunological responses to pregnancy between cows and heifers. We observed that cows had greater expression of proinflammatory cytokines and molecules associated with leukocyte migration and phagocytosis compared to heifers. Moreover, animals that failed to become pregnant showed altered expression of anti-inflammatory molecules. Overall, these findings support the hypothesis that early pregnancy signaling alters the proportions and functions of peripheral blood immune cells and differences between cows and heifers may yield insight into the reduced fertility of mature lactating dairy cows.

**Interpretative summary:** In dairy farming, 35-40% of high-yielding cows and 55-65% of nonlactating heifers conceive after their initial service. The immune system plays a crucial role in protecting cows from diseases and facilitating pregnancy establishment. This study investigates immune adaptation in bovine pregnancy by analyzing changes in immune cell populations and the expression of immunoregulatory molecules in the blood of heifers and lactating cows during early pregnancy. The findings support the hypothesis that early pregnancy signaling alters the proportions and functions of peripheral blood immune cells, offering insights into reduced fertility in mature lactating dairy cows.

## Introduction

Despite ongoing improvements in livestock husbandry and genetics, suboptimal conception rates in cattle still impair the economics, management, and sustainability of dairy farms. Most pregnancy losses occur within the first month after fertilization (Spencer, 2013; Diskin et al., 2016). During this period of early pregnancy, the bovine embryo secretes interferon tau (IFNT), a type I interferon that maintains a functional corpus luteum (CL) and progesterone secretion (Bazer et al., 1997). Further, the semi-allogeneic embryo also needs to avoid immune attack and facilitate endometrial remodeling for placentation (Pedersen et al., 2017; Ott, 2019). Thus, knowledge about changes in immune cells in the endometrium and peripheral blood as well as the immunoregulatory role of hormones such as IFNT and progesterone can improve understanding about the function of the immune system in pregnancy (Ott, 2019; Solano, 2019).

Relatively little is known about changes in immune cells in the uterus and peripheral blood during early pregnancy in cattle. In the endometrium, the number of NK cells and antigen presenting cells such as macrophages or dendritic cells was greater between day 8 to 20 of pregnancy compared to nonpregnant heifers (Oliveira and Hansen, 2008; Oliveira et al., 2013; Kamat et al., 2016; Vasudevan et al., 2017). CD4 T cells and B cells are present in the bovine endometrium around day 16, but the number of these cells did not differ between the estrous cycle and pregnancy (Leung et al., 2000). Expression of molecules such as indoleamine 2,3-dioxygenase 1 (IDO1), interleukin 10 (IL10), cytotoxic T-lymphocyte associated protein 4 (CTLA4), GATA binding protein 3 (GATA3), lymphocyte activating 3 (LAG3), and transforming growth factor beta (TGFB) also increased in the bovine endometrium during early pregnancy (Mansouri-Attia et al., 2012; Oliveira et al., 2013; Kamat et al., 2016; Vasudevan et al., 2017) suggesting the presence of local mechanisms to induce immune tolerance toward the conceptus (Ott, 2019).

In peripheral blood leukocytes (PBL), the percentage of monocytes was greater on day 17 in pregnant versus cyclic heifers (Kamat et al., 2016). Expression of the anti-inflammatory cytokines, IL10, interleukin 4 (IL4), TGFB, and the enzyme IDO1 was also greater in blood leukocytes of pregnant versus cyclic cows (Yang et al., 2016; Mohapatra et al., 2020). The authors also observed that transcript abundance for interleukin 1 (IL1), interleukin 6 (IL6), interleukin 8 (IL8) and IL10 were greater in the endometrium compared with peripheral blood mononuclear cells (PBMC) (Rutigliano et al., 2022). Clearly, the establishment of pregnancy involves regulation of immune cell types to direct migration, proliferation, apoptosis, and cytokine secretion in the uterus and circulation. However, there is still much that remains unexplained regarding immune adaptations during successful and failed pregnancies.

At breeding, dairy heifers are nulliparous, approximately 13 months of age, and have an anabolic metabolism. On the other hand, cows are lactating, multiparous, 50-80 days postpartum, and often experience effects of postpartum disease and catabolic metabolism to support energetic demands of lactation. Moreover, only 34% of high-yielding dairy cows and 54% of heifers and low-yielding cows calve after their first insemination (Diskin et al., 2016). Because immune cells are important for establishment of pregnancy and are sensitive to aging, nutrition, stress, and immune challenging events, we assessed whether heifers and cows differed in their immune responses during early pregnancy, perhaps contributing to disparities in fertility. Thus, we developed a series of experiments to characterize the changes in immune cell populations and expression of immunoregulatory molecules over days of the estrous cycle and early pregnancy in heifers and cows separately. Our first hypothesis was that the percentages of immune cell types and expression of immunoregulatory molecules in PBL change during early pregnancy compared to the estrous cycle. Subsequently, we compared the most striking immunological differences between cows and heifers during a critical period of early pregnancy where most embryo loss occurs. We hypothesized that cows have an elevated proinflammatory immune status compared to heifers that may relate to their reduced fertility.

## Material and Methods

### Ethics

This research was conducted with cattle from the Dairy Barn of Pennsylvania State University. Animal handling and experimental procedures were performed in accordance with the Guide for the Care and Use of Agricultural Animals in Research and Teaching and were approved by the Pennsylvania State University Institutional Animal Care and Use Committee (protocol #PRAMS201747548).

### Experiment 1

The aim of this experiment was to broadly characterize and compare changes in immune cell populations and expression of immunoregulatory molecules of heifers during different days of the estrous cycle and pregnancy. From 12/2018 to 04/2019, Holstein dairy heifers (N = 7, nulliparous, 13-14 months of age) were synchronized to estrus with an intramuscular injection of prostaglandin F2 alpha (PGF2A) *analog* (Estrumate, 500 μg cloprostenol sodium, Merck Animal Health) and monitored for estrus (day 0). Blood was collected between 8-10 am (during feeding) from the tail vein into K3 vacuette tubes (Greiner Bio-one) containing ethylenediaminetetraacetic acid (EDTA) on days 14, 17 and 20 of the estrous cycle. Later, the same animals were synchronized to estrus and inseminated, followed by blood collection on days 14, 17, 20 and 23 of pregnancy. Blood samples were used for PBL isolation and hormone assays. Pregnancy was confirmed on day 30 after insemination by transrectal ultrasonography. Two inseminated heifers became pregnant after the second service while the others carried first-service pregnancies. All pregnant animals delivered healthy calves.

### Experiment 2

This experiment had the same objective as in experiment 1, but it was conducted using lactating cows. From 06/2020 to 09/2020, Holstein dairy cows (first service, 60-70 days in milk, 2nd-5th parity) were synchronized to estrus via intramuscular injections of PGF2A (Lutalyse, 25 mg dinoprost tromethanmine, Zoetis) and gonadotropin releasing hormone (GnRH) (Factrel, 100 μg gonadorelin, Zoetis) following the PG-3-G Ovsynch (Peters and Pursley, 2002). Cows were assigned randomly to be either inseminated (N = 8) or not inseminated (N = 6) with blood collection on the same days as in experiment 1. Pregnancy was confirmed on day 30 after insemination using transrectal ultrasonography. All pregnant animals delivered healthy calves.

### Experiment 3

Based on results from experiment 1 and 2, a subset of immune molecules was selected to directly compare expression between parities (heifers vs cows) and reproductive statuses (cyclic, pregnant, and bred-nonpregnant). From 06/2021 to 10/2021, heifers and cows were synchronized to estrus weekly as described in experiment 1 and 2, using PGF2A (Estroplan, 500 μg cloprostenol sodium, Parnell) and GnRH (Gonabreed, 100 μg gonadorelin acetate, Parnell). Animals were assigned randomly to be either inseminated or not inseminated to generate the following treatment groups: cyclic heifers (N = 10), pregnant heifers (N = 8), cyclic cows (N = 7), and pregnant cows (N = 9). Blood was collected as described in experiment 1, but only on days 17 and 18 of the estrous cycle and pregnancy. Cows and heifers were recruited and synchronized simultaneously so days of blood collection and flow cytometric analysis included samples of each parity and reproductive status. Concentration of progesterone in plasma from days 17 and 18 was used to confirm the presence of a functional CL in all animals while blood from day 18 was used for PBL isolation and downstream immune cell analysis. We selected day 18 for sample collection to allow sufficient time for conceptus signaling to affect peripheral immune cells but also ensure that the CL would still be functional in cyclic animals. Pregnancy was detected via transrectal ultrasonography on day 30 following insemination and heifers and cows confirmed nonpregnant were grouped in the bred-nonpregnant (BNP) status. All pregnant animals delivered healthy calves.

### Leukocyte isolation and flow cytometry

Total PBL were isolated from approximately 20 mL of blood following the protocol described in Kamat et al. (2016). For flow cytometry analysis of intracellular proteins, 0.25 million cells were added to duplicate wells of a round bottom 96-well plate (Corning Falcon) and fixed with cytofix/cytoperm reagent (BD Biosciences) following the manufacturer’s protocol. For the remaining surface proteins, 0.5 million cells were added to each well of a round bottom 96-well plate. Antibody incubation followed the protocol described in Kamat et al. (2016), using: (1) 15% bovine serum albumin (BSA) in 10% Perm/Wash during antibody labeling of intracellular proteins and 10% Perm/Wash during cell washes and (2) Phosphate-buffered saline with ethylenediaminetetraacetic acid (PBS-EDTA) during antibody labeling of surface proteins and cell washes. At the end of all antibody incubations, cells were suspended in PBS-EDTA buffer and analyzed on the Guava EasyCyte flow cytometer using a 488 nm blue laser for fluorescein isothiocyanate (FITC) fluorophore, setting voltages based on control wells (cells without antibody, with only secondary antibody, and with isotype control), and counting 30,000 events/well. The data obtained were analyzed using FlowJo software (FlowJo LLC). Flow cytometry gating strategy is described in supplementary data.

Antibodies and concentrations used during flow cytometry analyses are described in Table 1. The molecular targets include IDO1, aryl hydrocarbon receptor (AHR), peroxisome proliferator activated receptor gamma (PPARG) and the surface molecules CD8B, CD14, CD47, CD3 epsilon subunit of T-cell receptor complex (CD3E), natural cytotoxicity triggering receptor 1 (NCR1; also known as NKP46 or CD335), complement C3d receptor 2 (CR2; also known as CD21), integrin subunit alpha M (ITGAM; also known as CD11B), integrin subunit alpha X (ITGAX; also known as CD11C), signal regulatory protein alpha (SIRPA), and MM20A granulocyte epitope. It is worth noting that MM20A is an antibody specific for bovine granulocytes, but the protein targeted by this antibody has not been described. Considering that ∼80% of blood granulocytes are neutrophils, we used MM20A to evaluate changes in neutrophils in the blood of dairy cattle.

**Table 1:**
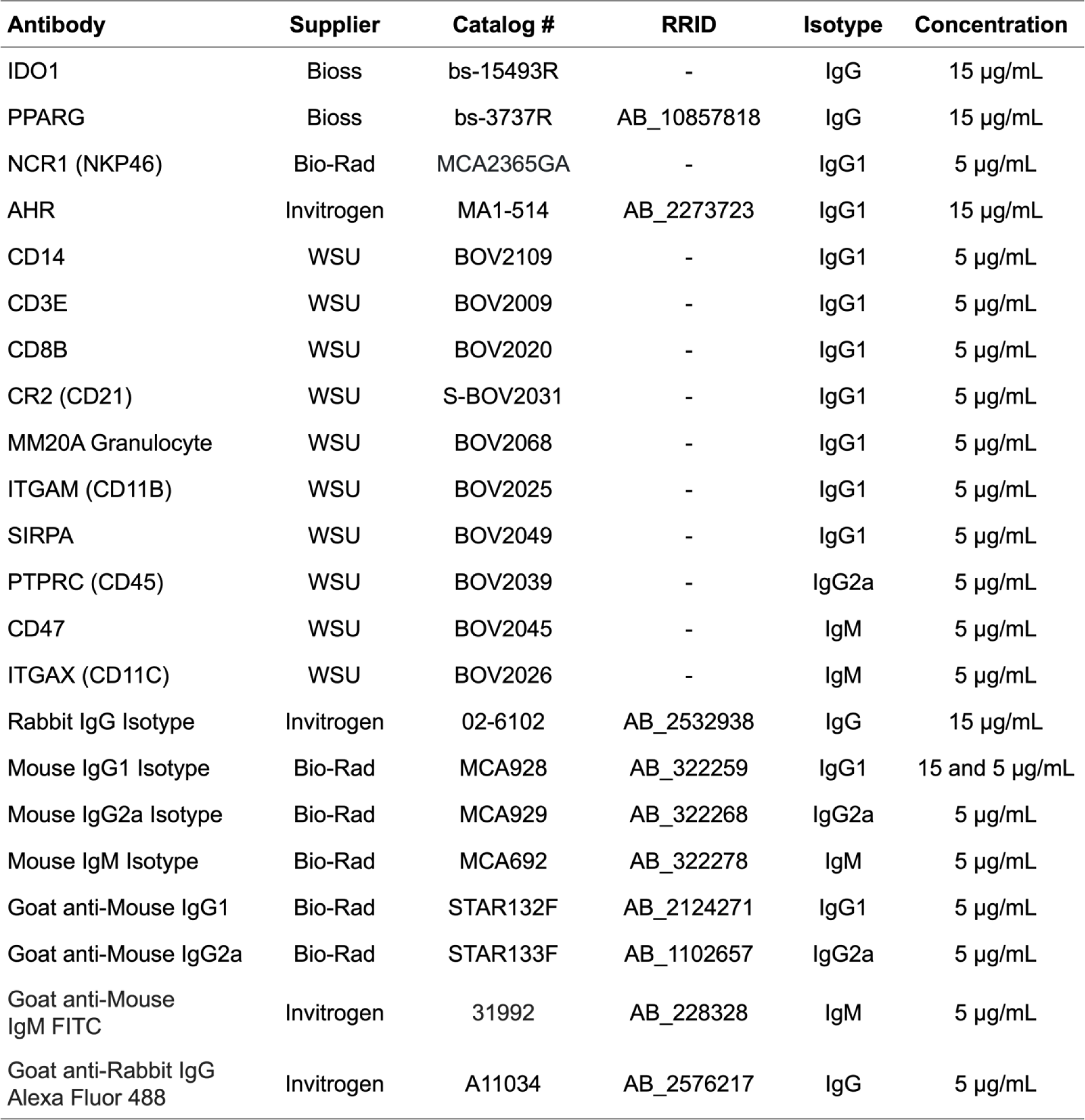
Antibodies used in flow cytometry labeling. Legend: Washington State University (WSU); Research Resource Identifier (RRID).

### Progesterone ELISA

Concentration of progesterone in plasma was determined for all animals in Experiment 3 using an ELISA protocol previously validated for cattle (Hughes et al., 2021). The inter-assay coefficients of variation (CV) across 3 plates were 3.93% and 1.70% based on the same high and low progesterone standard controls. The intra-assay CV for plates 1, 2 and 3 were 5.13%, 6.20%, and 6.90%, respectively.

### RT-qPCR

Total cellular RNA was isolated from approximately 10 million cells using 500 µL of TRIzol reagent (Life Technologies). Quantity and quality of RNA was assessed using the Experion electrophoresis system (Biorad) according to the manufacturer’s protocol. All samples showed RNA quality indicator (RQI) > 8. Primer sets (Table 2) were validated and used in reverse transcription quantitative polymerase chain reaction (RT-qPCR) analysis. Standard curves showed efficiency between 83% to 99%, and samples with cycle threshold below detection limit of the standard curve were removed from data analysis. Ribosomal protein L19 (RPL19) was selected as the reference gene and its cycle threshold (Ct) values were not affected by treatment. Gene expression was analyzed for IDO1, AHR, PPARG, CD3E, IL10, IL4, IL6, interferon gamma (IFNG), fatty acid binding protein 4 (FABP4), and cytochrome P450 family 1 subfamily A member 2 (CYP1A2).

**Table 2:**
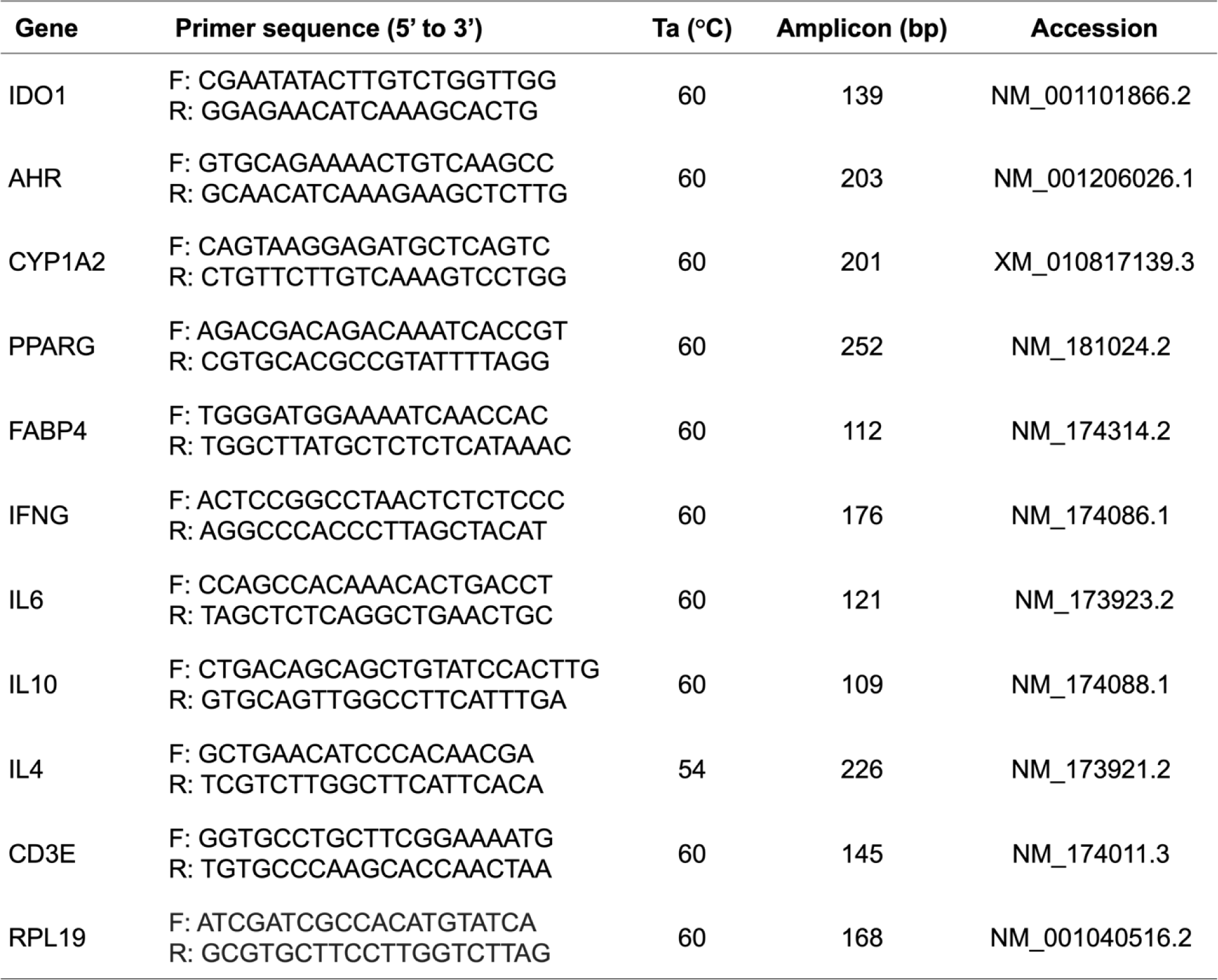
Primer used in RT-qPCR analysis. Assay was validated to confirm the annealing temperature (Ta), amplicon size and sequence of each primer set.

### Statistical analysis

Statistical analyses were performed using Mixed model procedures of SAS (v 9.4; Statistical Analysis System Institute). Probability values < 0.05, and 0.05 < p < 0.10 were considered statistically significant and a tendency for significance, respectively. Normal distribution of data was assessed visually and using Shapiro Wilk test. Data sets lacking normal distribution were log-normal transformed and rechecked to confirm normality. Outliers were removed if studentized residuals were greater than the absolute value of 3. In experiment 1 and 2, two analysis of variance (ANOVA) models were conducted. The first model fitted the effects of status (cyclic vs pregnant), day (14, 17, 20), and their interaction on flow cytometry parameters of percent positive cells and mean fluorescence intensity (MFI). The second model was performed to predict the relationship between flow cytometry parameters and days of pregnancy (14, 17, 20, and 23) using a polynomial regression analysis. Both models included the variable, animal, as a repeated effect for experiment 1 and as a random effect for experiment 2. Moreover, because data collection was done on different days and for many weeks, the effect of assay run was also tested and included in the model to adjust the least squares means when significant.

In experiment 3, due to differences in progesterone concentration in plasma of cyclic heifers, an ANOVA was conducted first to assess whether progesterone concentration was a source of variation among flow cytometry parameters. When the effect of progesterone concentration was significant, cyclic heifers with regressed luteal status (N = 7) were removed from the second statistical analysis. The second ANOVA model fitted the effects of parity (heifers vs cows), status (cyclic vs pregnant vs bred-nonpregnant), and parity*status interaction on flow cytometry parameters. All statistical models included the variable animal as a random effect. The effect of assay run was also tested and included in the analysis when significant.

Graphs were generated using GraphPad (v. 9.3; GraphPad Software Inc.). Flow cytometry results were plotted as least squares mean (LSM) ± standard error mean (SEM) while the LSM of transformed data were back transformed by raising 10 to the power of LSM and plotted with 95% confidence interval (CI). For RT-qPCR data in experiment 3, the 2^−ΔΔCt^ values were calculated relative to day 18 of estrous cycle in heifers and used during statistical analyses for the main effect of parity, status, and parity*status interaction. The data was graphed as LSM ± SEM. When parity was a significant source of variation, a second statistical analysis was performed to assess the effect of status within parity, also using 2^−ΔΔCt^ values. Following a significant F-test, differences in LSM between groups in all analysis were determined by Tukey’s multiple comparisons adjustment.

## Results

### Experiment 1

Among the leukocyte populations analyzed in heifers, the percentage of monocytes (CD14^+^) tended to be greater during pregnancy than the estrous cycle (Status: p = 0.06; Figure 1A). The percentage of T cells (CD3E^+^) in pregnant heifers differed from that in cyclic heifers at day 14, 17 and 20, but there was no consistent pattern. (Status*Day: p = 0.06; Figure 1B). Moreover, the percentage of T cells (CD3E^+^: p = 0.04), cytotoxic T cells (CD8B^+^: p < 0.01), monocytes (CD14^+^: p < 0.01), B cells (CR2^+^: p = 0.07), and NK cells (NCR1^+^: p < 0.01) increased linearly over days of pregnancy in heifers (Figure 1C), although the increase was small. The abundance/cell of several proteins associated with immune cell type or function was greater or tended to be greater in pregnant compared to cyclic heifers. These proteins include surface receptors such as CD8B (Status: p = 0.05; Figure 1D), CR2 (Status: p = 0.02; Figure 1E), NCR1 (Status: p = 0.07; Figure 1F), CD14 (Status: p = 0.03; Figure 1G), MM20A granulocyte epitope (Status: p = 0.03; Figure 1H), CD47 (Status: p = 0.01; Figure 1I), and SIRPA (Status: p = 0.01; Figure 1J). Moreover, from day 14 to day 23, expression of MM20A granulocyte antigen (Day: p = 0.08), NCR1 (Day: p = 0.04), CD14 (Day: p = 0.02), and IDO1 (Day: p = 0.02) decreased or tended to decrease in pregnant heifers (Figure 1K). Immune parameters measured that did not exhibit changes in experiment 1 are shown in supplementary data.

**Figure 1:**
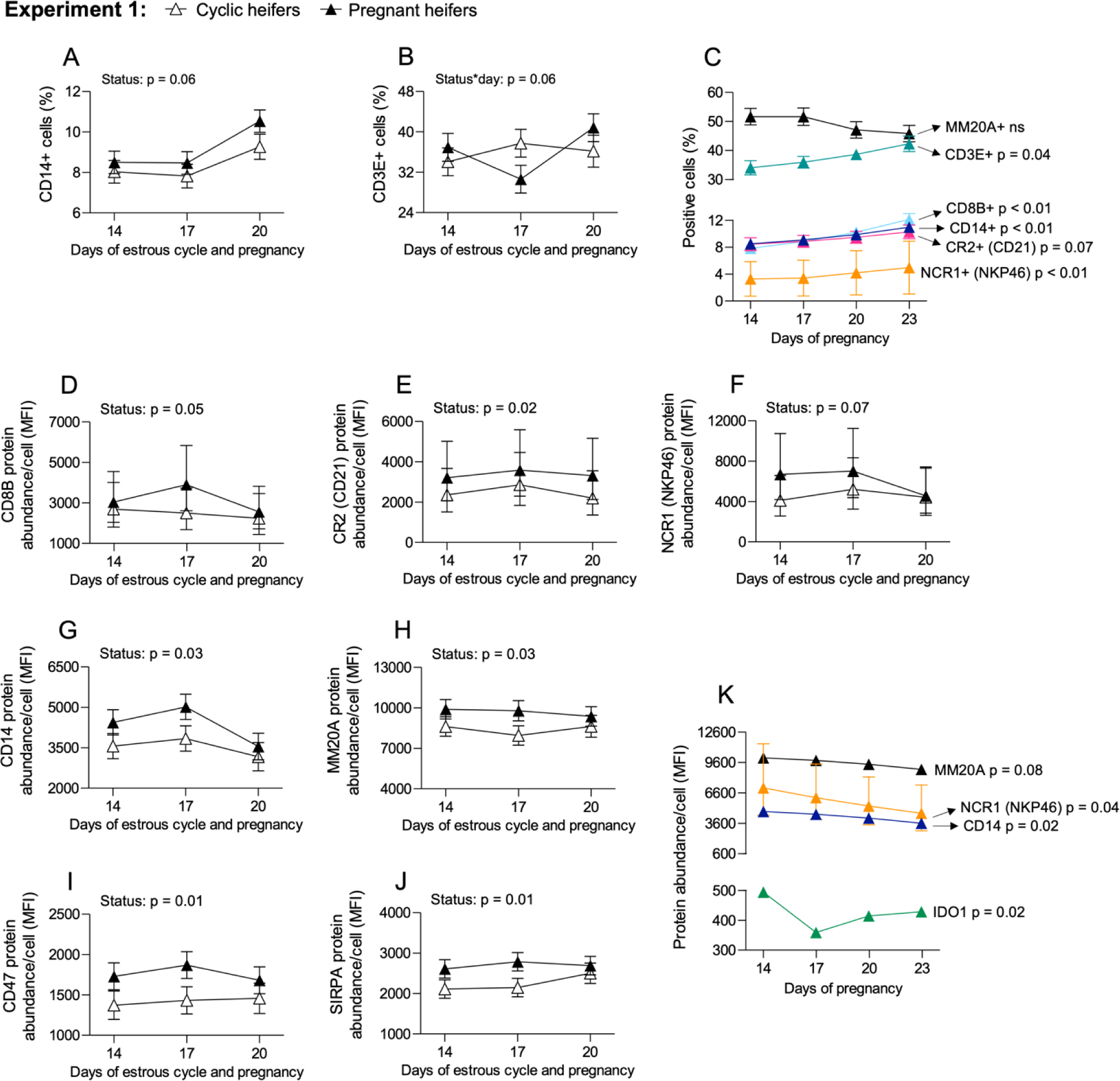
Percentage of leukocytes and protein expression of immunoregulatory molecules in heifers from experiment 1. Changes in the percentage of (A) monocytes and (B) total T cells in cyclic (white triangle, N = 7) versus pregnant (black triangle, N = 7) animals. (C) Changes in percentage of leukocyte subsets over days of early pregnancy. Protein expression/cell of (D) CD8B, (E) CR2, (F) NCR1, (G) CD14, (H) MM20A, (I) CD47, and (J) SIRPA in cyclic compared to pregnant animals. (K) Protein expression/cell of immunoregulatory molecules over days of pregnancy.

### Experiment 2

The percentages of cytotoxic T cells (CD8B^+^, Status*Day: p = 0.03; Figure 2A) and NK cells (NCR1^+^, Status*Day: p = 0.02; Figure 2B) were less in pregnant compared to cyclic cows on day 17 and 20. Over days of pregnancy, the percentage of T cells (CD3E^+^: p = 0.08) and monocytes (CD14^+^: p = 0.05) tended to decrease from day 14 to 17 while the percentage of NK cells (NCR1^+^: p < 0.01) decreased over days of pregnancy (Figure 2C). Pregnant cows had greater protein expression/cell of only two immune molecules compared to cyclic cows: MM20A granulocyte epitope (Status: p = 0.04; Figure 2D) and CD47 (Status: p = 0.01; Figure 2E). A decrease in protein expression of molecules involved in the function of phagocytes such as MM20A granulocyte epitope (Day: p < 0.01), SIRPA (Day: p < 0.01), CD47 (Day: p = 0.02), ITGAX (Day: p < 0.01), AHR (Day: p = 0.05), and PPARG (Day: p = 0.01) was also observed over days of pregnancy in cows (Figure 2F). Immune parameters measured that were not affected by status or day in experiment 2 are shown in supplementary data.

**Figure 2:**
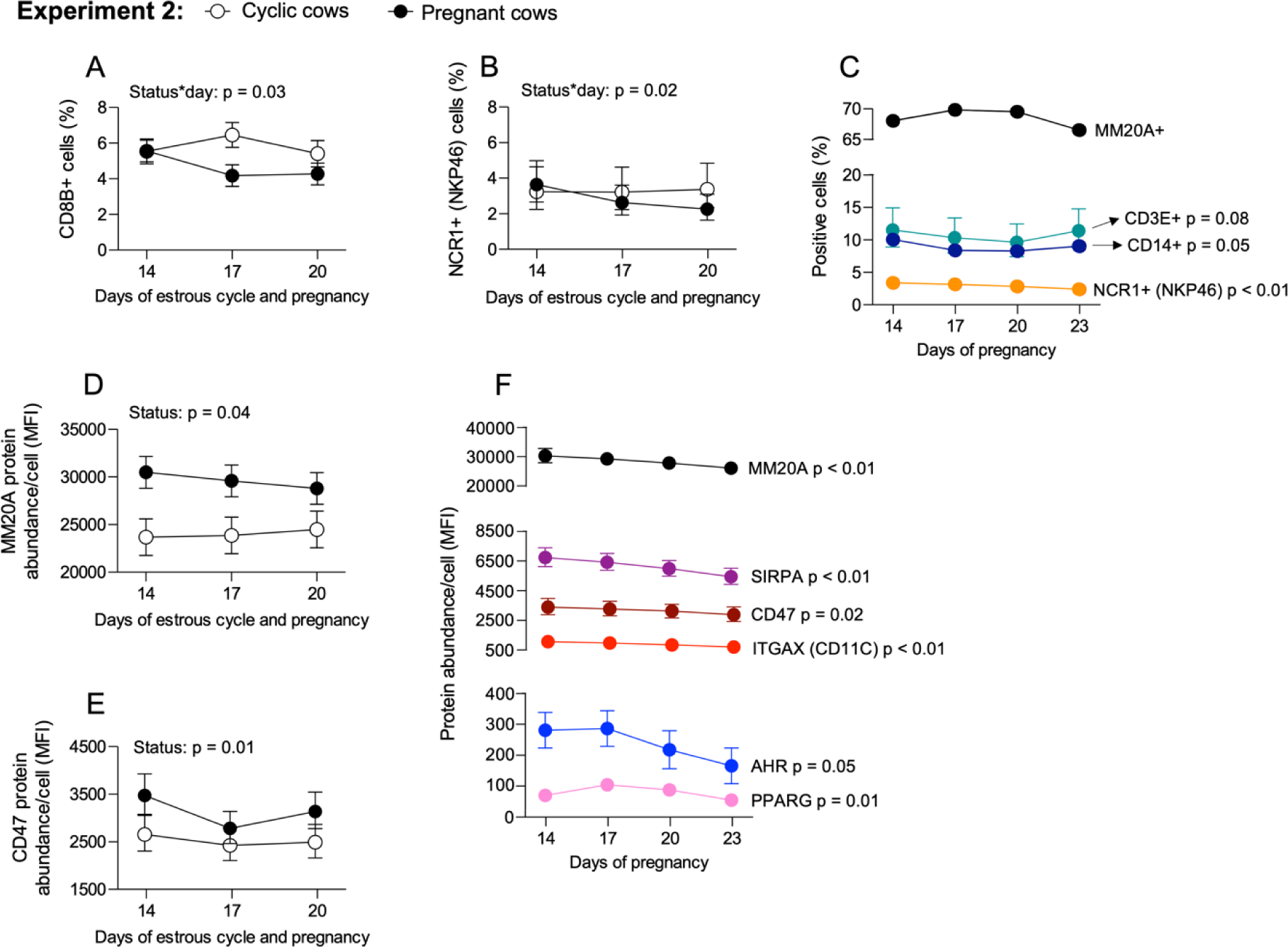
Percentage of leukocytes and protein expression of immunoregulatory molecules in cows from experiment 2. Percentage of (A) CD8B+ T cells and (B) NK cells in cyclic (white circle, N = 6) versus pregnant (black circle, N = 8) animals. (C) Changes in percentage of leukocyte subsets over days of early pregnancy. Protein expression/cell of (D) MM20A and (E) CD47. (F) Protein expression/cell of immunoregulatory molecules over days of pregnancy.

### Experiment 3

Analysis of plasma progesterone showed that 6/10 cyclic heifers had a regressed CL (low plasma progesterone) by day 18 whereas the CL was maintained in pregnant animals and in cyclic cows (Figure 3). Therefore, we included plasma progesterone in the statistical model. Comparing effects of parity on PBL proportion and phenotype, MM20A^+^ neutrophils (p = < 0.01; Figure 4A) SIRPA^+^ (p < 0.01; Figure 4B), ITGAM^+^ (p < 0.01; Figure 4C), and ITGAX^+^ (p = 0.04; Figure 4D) were 20-30% greater in cows than in heifers. In addition, lower expression of CD3E mRNA in PBL of cows compared to heifers (p = 0.01; Figure 4E) may indicate differences in proportion of T cells between parities. Cows had greater IFNG (p = 0.03; Figure 4F) and IL6 (p = 0.01; Figure 4G) mRNA abundance in PBL than heifers, but no difference in TNF mRNA was observed between parities (data not shown). AHR protein abundance/cell was also greater in cows than heifers (p < 0.01; Figure 4H), although AHR mRNA was not different (Figure 4I). Consistent with AHR protein expression, CYP1A2 mRNA (gene regulated by AHR activation) tended to be greater in cows than heifers (p = 0.08; Figure 4J). When parity was a significant source of variation for a given parameter, we tested the effects of status in heifers and cows separately. Only cows showed differences in immune molecule expression related to their reproductive status. Pregnant cows tended to have a greater percentage of cells positive for SIRPA (p = 0.08; Figure 4B) and ITGAM (p = 0.08; Figure 4C) than cyclic cows, but no difference between pregnant and bred-nonpregnant animals was observed. PBL of bred-nonpregnant cows had less CD3E mRNA than cyclic cows but were similar to pregnant cows (p = 0.01; Figure 4E).

**Figure 3:**
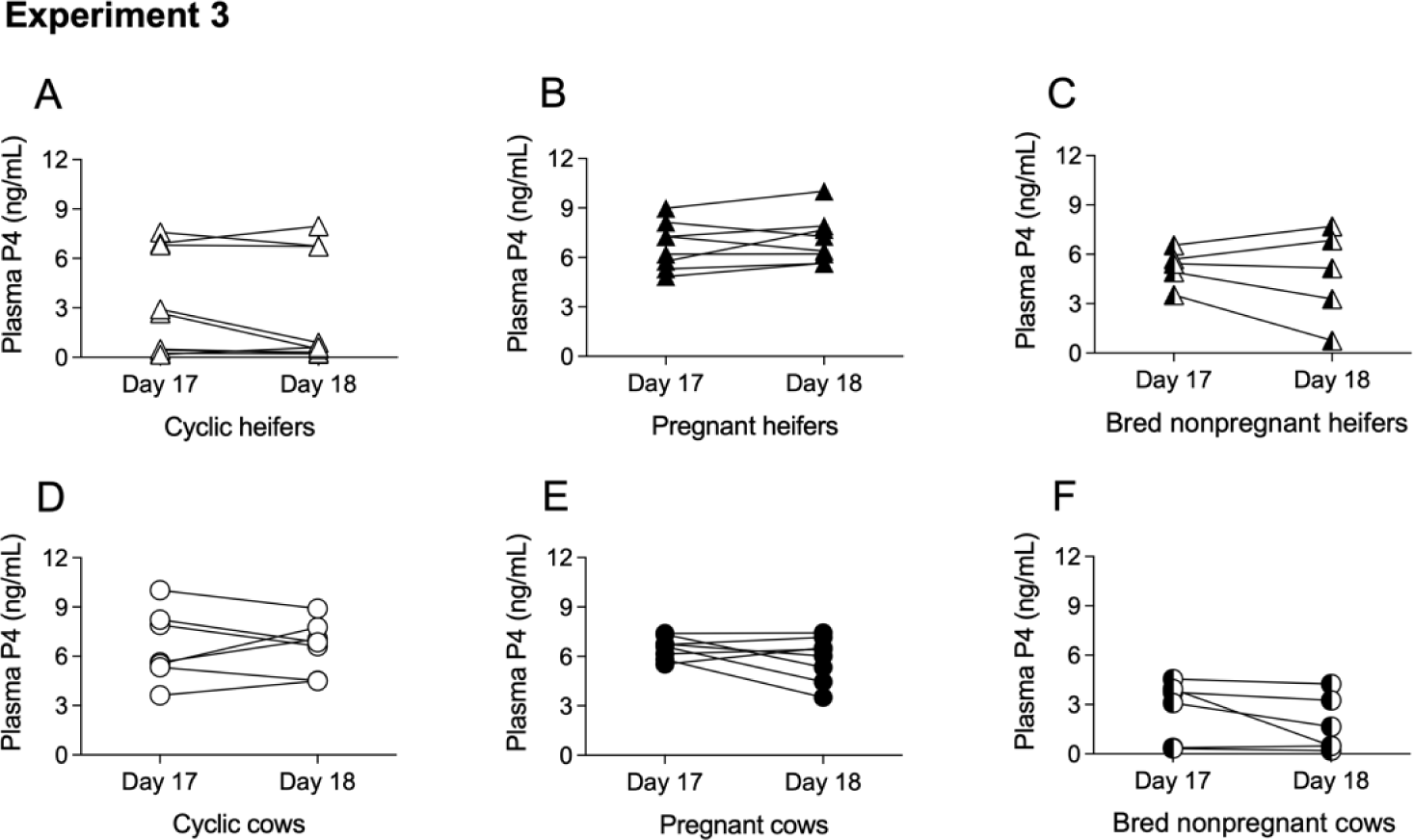
Plasma progesterone (P4) in cyclic, pregnant, and bred-nonpregnant (A-C) heifers and (D-F) cows from experiment 3. Note that 7 of 10 heifers showed low P4 or regressing/regressed corpus luteum prior day 18 of estrous cycle.

**Figure 4:**
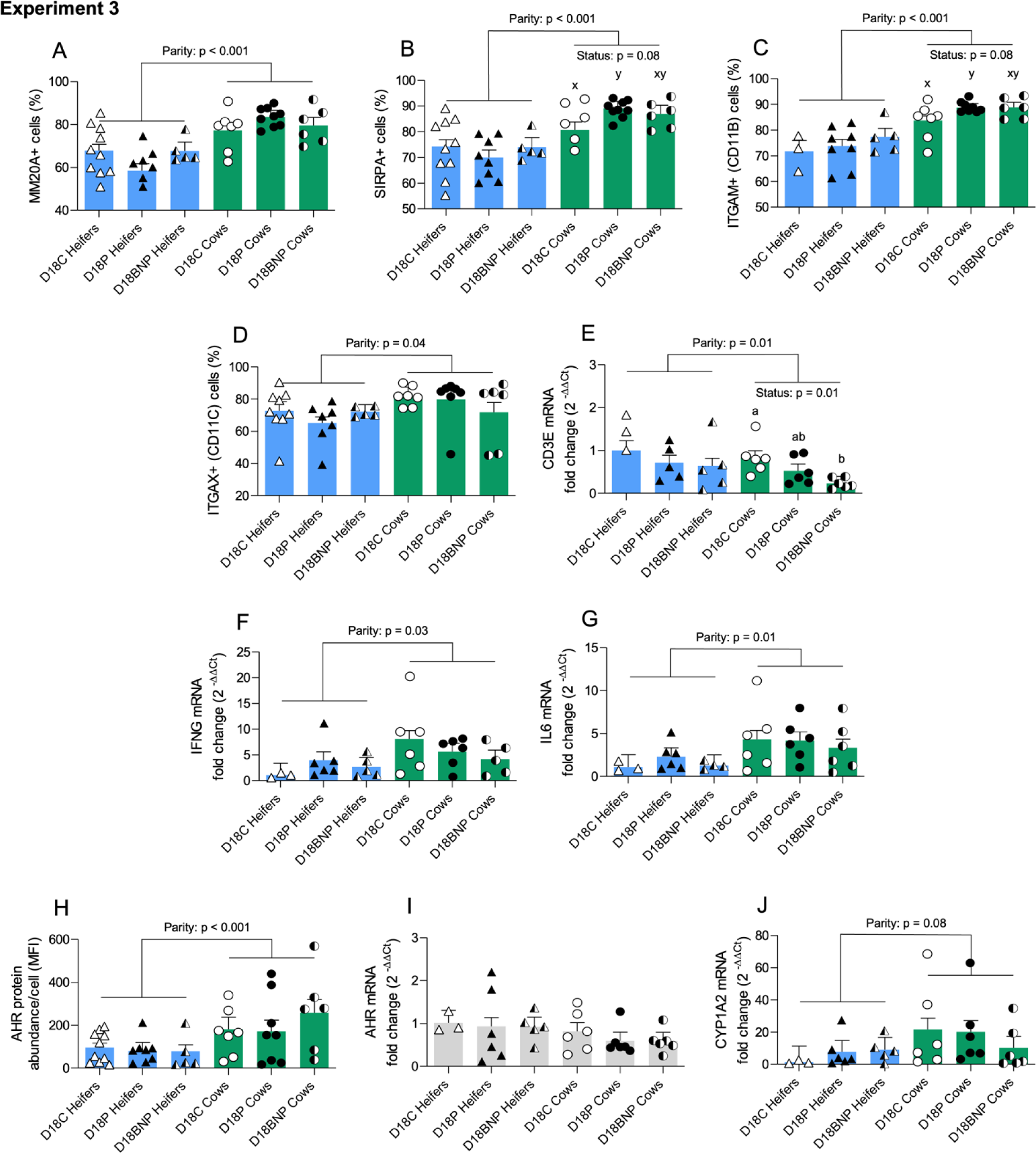
Percentage of leukocytes and immunoregulatory molecules differentially expressed between heifers and cows from experiment 3. Changes in percentage of (A) neutrophils, (B) SIRPA^+^ cells, (C) ITGAM^+^ cells, and (D) ITGAX^+^ cells. mRNA fold change of (E) CD3E, (F) IFNG, and (G) IL6. Protein expression/cell of (H) AHR and mRNA expression of (I) AHR and its target gene (J) CYP1A2. The mRNA fold change was calculated relative to cyclic heifers. When parity was a significant source of variation, the effect of status was assessed within parity using values of mRNA fold change relative to their respective cyclic animals. Legends: day 18 of estrous cyclic (D18C); day 18 of pregnancy (D18P); day 18 of bred-nonpregnant (D18BNP); a, b when p < 0.05 and x, y when 0.5 > p < 0.10 for effect of status.

Animals that failed to become pregnant exhibited altered expression of some immune molecules. IL10 mRNA in PBL of bred-nonpregnant heifers were greater than in bred-nonpregnant cows (p = 0.03; Figure 5A) and IL4 mRNA tended to be greater in pregnant heifers and cows than in cyclic or bred-nonpregnant animals (p = 0.10; Figure 5B). The percentage of PBL positive for PPARG was less in bred-nonpregnant heifers and cows than the other statuses (p = 0.01; Figure 5C), however, protein abundance/cell of this molecule tended to be greater in bred-nonpregnant cows (p = 0.01; Figure 5D). PPARG mRNA (p = 0.03; Figure 5E) as well as FABP4 mRNA (p = 0.03; Figure 5F), a gene regulated by PPARG activation, were greater in cows than heifers. The percentage of cells positive for IDO1 was less in PBL of bred-nonpregnant animals (p < 0.01; Figure 5G), but no changes were observed in IDO1 protein abundance/cell or mRNA (Figure 5H-I).

**Figure 5:**
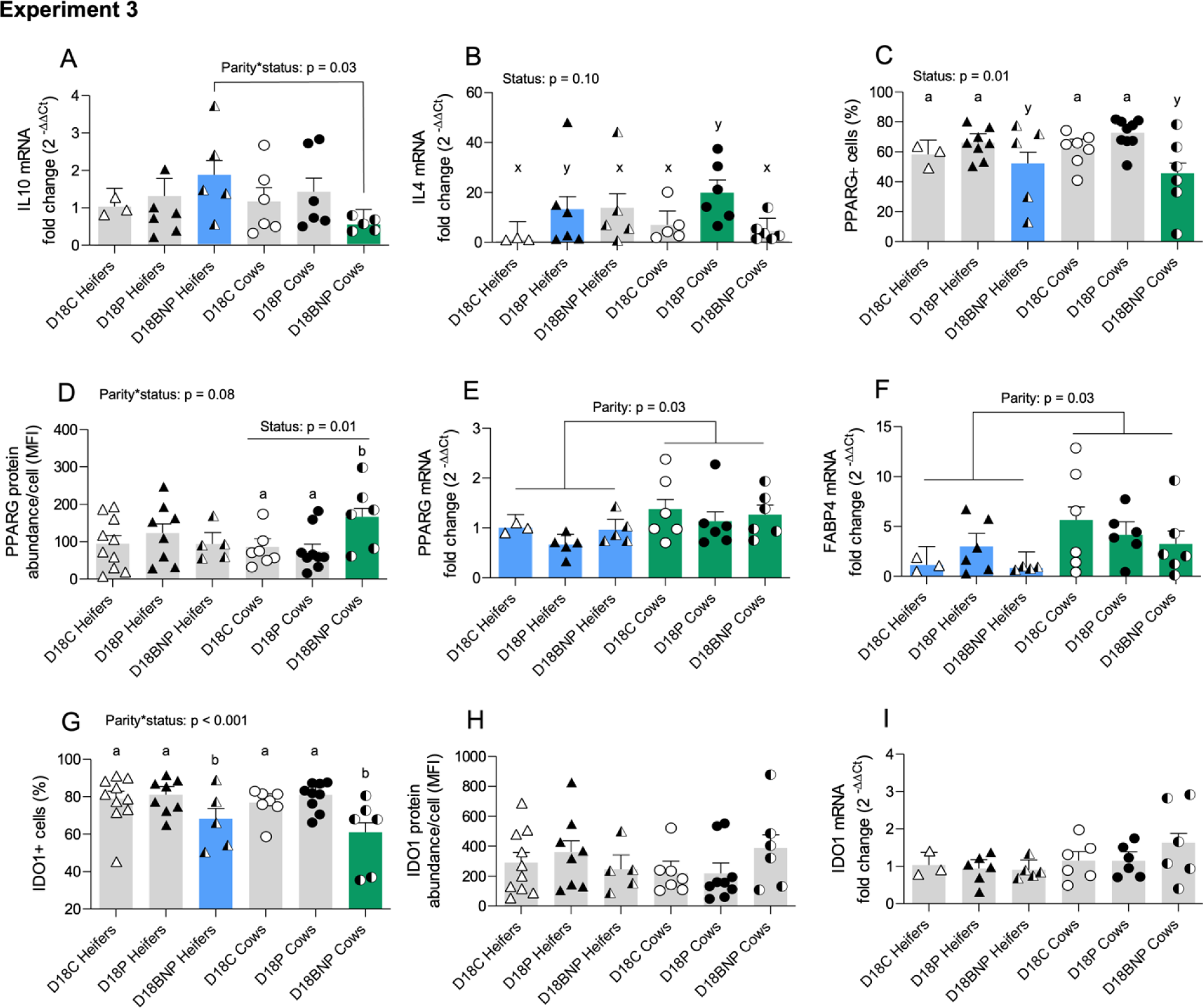
Percentage of leukocytes and immunoregulatory molecules differentially expressed in bred-nonpregnant animals from experiment 3. mRNA fold change of (A) IL10, and (B) IL Percentage of (C) PPARG^+^ cells and (D) protein abundance/cell and (E) mRNA expression of PPARG and its target gene (F) FABP Proportion of (G) IDO1^+^ cells and (H) protein abundance/cell and (I) mRNA expression of IDO1. The mRNA fold change was calculated relative to cyclic heifers. When parity was a significant source of variation, the effect of status was assessed within parity using values of mRNA fold change relative to their respective cyclic animals. Legends: day 18 of estrous cyclic (D18C); day 18 of pregnancy (D18P); day 18 of bred-nonpregnant (D18BNP); a, b when p < 0.05 and x, y when 0.5 > p < 0.10 for effect of status.

## Discussion

The substantial difference in conception rates between cows and heifers creates inefficiencies and economic loss for dairy production. Because pregnancy is an immunologically unique phenomenon, the purpose of this study was to describe changes in circulating immune cells during early pregnancy of heifers and cows to better understand if immunological differences between these parities contributes to reduced fertility in cows. Although early pregnancy changed the phenotype of specific leukocyte subtypes in heifers and cows, our main findings showed that cows appear to have greater expression of proinflammatory cytokines and molecules associated with leukocyte migration and phagocytosis compared to heifers. Moreover, animals that failed to become pregnant showed altered expression of anti-inflammatory molecules. Overall, these findings support the hypothesis that immune responses to early pregnancy may differ in heifers and cows and is associated with reduced fertility of lactating dairy cows.

### Effect of pregnancy on the proportion of leukocyte subsets

Early pregnancy changed the proportion of specific, yet different, leukocyte subtypes in heifers and cows. In experiment 1, pregnant heifers had a greater percentage of monocytes and CD3+ T cells compared to cyclic heifers. This result agrees with our prior work showing a 2-fold greater percentage of CD14+ITGAX+ monocytes in the blood of day 17 pregnant compared to cyclic heifers (Kamat et al., 2016). An increase in the percentage of circulating monocytes was also seen in women and rats during the first (Luppi et al., 2002) and third trimester (Al-ofi et al., 2012; Melgert et al., 2012) of normal pregnancies. In cows, circulating CD68+ monocytes were reported to be greater in day-33 pregnant versus nonpregnant animals (Oliveira and Hansen, 2008). However, we did not observe differences in monocyte population between pregnant and cyclic cows in experiment 2, possibly because sampling occurred on early days of pregnancy. The percentage of T cells in pregnant heifers from experiment 1 was transiently less than cyclic heifers on day 17. This effect on the proportion of total T cells could be related to endocrine exposure to interferon during bovine pregnancy. High levels and/or prolonged interferon signaling have been reported to inhibit proliferation of T cells in chronic viral infection models (Le Saout et al., 2014; Cheng et al., 2017; Dagenais-Lussier et al., 2018).

In experiment 2, pregnant cows exhibited lower percentage of cytotoxic T and NK cells than cyclic cows. Others have reported that CD4+ and CD8+ lymphocytes remain unchanged in blood and endometrium of cows (Leung et al., 2000; Oliveira and Hansen, 2008) or increase in endometrium of heifers during early pregnancy (Vasudevan et al., 2017). Using an NK cell-specific reporter mouse and the parabiosis procedure, it was determined that during pregnancy, some uterine NK cells originated from the bloodstream and some from proliferation of tissue resident cells (Sojka et al., 2018). Thus, it is possible that the decrease in cytotoxic T and NK cells in pregnant cows from experiment 2 may result from migration of these leukocytes from blood into the uterus.

Considering only the results from pregnant animals, a temporal increase in percentage of CD3E+, CD8B+, CD14+, CR2+, and NCR1+ cells in heifers from experiment 1 was consistent and was accompanied by a non-significant decrease in frequency of MM20A+ neutrophils. Similarly, changes in the percentage of neutrophils seemed to affect the proportion of CD3E+, CD14+, NCR1+ in cows from experiment 2. However, interpretation of percentage data in flow cytometry requires consideration of the overall sample composition and the presence of other cell subsets. For example, changes in percentage of lowly abundant leukocyte subsets may result from alterations in the frequency of an abundant leukocyte population such as neutrophils. Moreover, the proportion of leukocytes can also be changed by processes of cell differentiation, mobilization of bone marrow-derived precursor cells, proliferation, apoptosis, demargination, and diapedesis (Blumenreich, 1990; Mohapatra et al., 2020).

### Effect of pregnancy on the protein expression of immune molecules

Pregnant heifers from experiment 1 showed greater protein abundance/cell than cyclic heifers for multiple immune function molecules. On the other hand, pregnant cows from experiment 2 only showed greater protein expression of MM20A granulocyte epitope and CD47 compared to cyclic animals. These differences may relate to the experimental design for each experiment. The repeated measure design in experiment 1 can reduce inter-animal biological variation, whereas selecting cows at random in experiment 2 may increase data variability and mask pregnancy effects on protein expression. It is also possible that pregnant heifers show greater protein abundance/cell of immune molecules because of exposure to IFNT or other embryonic antigens for the first time. For example, the expression of interferon stimulated genes on day 18 of pregnancy was shown to be greater in PBL of nulliparous heifers than in multiparous cows (Green et al., 2010).

Interferons are potent activators of innate immune responses and upregulate numerous surface molecules that increase immune sensing and elimination of pathogens (Belardelli, 1995). Interferon alpha (IFNA) and IFNG increase expression of NCR1 (Bozzano et al., 2011), ITGAX (Barbalat et al., 2009), CD47 (Ye et al., 2021), SIRPA (De Almeida et al., 2012; Myers et al., 2019), CD14 (Lee and Sullivan, 2001) and the number of CD8+ T cells (Whitmire et al., 2005), all molecules that were greater in pregnant versus cyclic heifers from experiment 1. Interferon signaling can also elicit immunological memory (Jaitin and Schreiber, 2007; Kamada et al., 2018) and regulate proliferation and apoptosis of memory leukocytes (Zhang et al., 1998; Bahl et al., 2010; Huber and David Farrar, 2011; Welsh et al., 2012). However, development of immunological memory to antigens of a first pregnancy and its effect in subsequent gestations is poorly understood and has not been investigated in cattle but could play a role in immune response during the establishment of pregnancy.

Considering the results from early pregnancy alone, the overall expression of immunoregulatory proteins decreased over time in both experiments, except for IDO1 in heifers from experiment 1 and AHR in cows from experiment 2. Heifers showed a ∼30% decrease in IDO1 protein in PBL day 14 and 17 of pregnancy. However, IDO1 mRNA was reported to be greater in PBL of cows from day 18 of pregnancy compared to the estrous cycle (Mohapatra et al., 2020). In the endometrium of heifers, IDO1 protein abundance was also greater on day 17 of pregnancy versus the estrous cycle (Groebner et al., 2011; Vasudevan et al., 2017). It is possible, even likely, that local and systemic immune responses to pregnancy differ between heifers and cows. Moreover, the differences in results reported in the literature may be due to discrepancies in animal group (heifers or cows), collected samples, and methods.

AHR protein expression/cell also decreased ∼40% between day 17 and 23 of pregnancy in cows and to the best of our knowledge, there are no publications about AHR expression in bovine PBL. AHR is a transcription factor that can be activated by metabolites downstream of IDO1 enzymatic activity (Gargaro et al., 2021). Effects of IDO1 via kynurenine-AHR signaling are proposed to be important for avoiding embryo rejection (Baban et al., 2004; Miwa et al., 2006) and inducing immunotolerance in antigen presenting cells (Mellor and Munn, 2004; Mezrich et al., 2010). Moreover, binding of DNA from apoptotic cells to toll-like receptor 9 (TLR9) also activates AHR in macrophages resulting in an increase in IL10, immune suppression and tolerance (Shinde et al., 2018). Interferons are apoptotic signals (Chawla-Sarkar et al., 2003) and considering the high levels of IFNT secreted during ∼4 weeks of early bovine pregnancy, decreased IDO1 and AHR protein in PBL may be a mechanism to promote immune activation instead of immunotolerance in circulation.

### Differences in PBL phenotype between heifers and cows

Studies comparing the composition of peripheral blood immune cells between cows and heifers are scarce. However, hematological evidence suggests that cows have similar proportions of lymphocytes (40-60%) and neutrophils (30-50%) (George et al., 2010; Herman et al., 2018) while heifers have greater proportion of lymphocytes (50-70%) than neutrophils (20-40%) (Ahmadi et al., 2006; Botezatu et al., 2014). Experiment 3 was designed to compare immunological differences between heifers and cows under the same experimental conditions. Results confirmed that cows have greater percentages of neutrophils, and myeloid cells positive for SIRPA and ITGAX than heifers. We also observed lower mRNA expression of CD3E in cows compared with heifers, agreeing with the findings in experiments 1 and 2 that suggested cows had a lower percentage of CD3E+ cells in PBL than heifers.

Neutrophils are the most abundant cell type of the innate immune system, often described as the first line of defense for clearing bacterial infection. However, neutrophils also assist with normal ovarian, uterine, gestational, and postpartum physiology in cattle (Alhussien and Dang, 2019). Around calving, high concentrations of cortisol induce neutrophilia (Burton et al., 2005; McDougall et al., 2017) important to facilitate placental expulsion and eliminate infection in the uterus and mammary gland (Moretti et al., 2016; Alhussien and Dang, 2019). After calving, cortisol concentration decreases, yet is higher in cows than heifers (Burnett et al., 2015). Thus, in relation to our results, postpartum lactating cows may have a greater percentage of circulating neutrophils and myeloid cells positive for SIRPΑ, ITGAM and ITGAX than nulliparous heifers due to effects of glucocorticoids on granulopoiesis, neutrophil longevity, and transmigration (Burton et al., 2005).

We also observed greater concentrations of mRNA for the proinflammatory cytokines, IFNG and IL6, in PBL of cows compared with heifers. IFNG is secreted by activated immune cells in response to pathogen associated molecular patterns (PAMPs), cytokines and type I interferons (Castro et al., 2018). Consequently, IFNG signaling activates transcription of proinflammatory genes that amplify the interferon response (Libermann and Baltimore, 1990). McLoughlin et al. (2003) showed that in a knockout mouse model, expression of IL6 in neutrophils depended on IFNG signaling and both cytokines regulate recruitment, apoptosis, and clearance of neutrophils during peritoneal inflammation. Cows also had greater protein expression/cell of AHR and a tendency for greater CYP1A2 mRNA, a target of AHR activation. AHR expression can be induced by IL6 signaling and AHR activation can upregulate transcription and synthesis of IL6 (Stobbe-Maicherski et al., 2013; Huang et al., 2022). Interestingly, AHR activation is involved with wasting syndrome (Girer et al., 2021) and obesity related fatty liver and inflammation (Pohjanvirta, 2017; Bock, 2021) and participates in leukocyte energy metabolism, neutrophil chemotaxis, and bacterial clearance (Partida-Sánchez et al., 2001; Bunaciu et al., 2015). Therefore, AHR may be an important regulator of leukocyte phenotype in the physiology of periparturient dairy cows.

When multiparous cows transition from pregnancy to a postpartum and lactating status, there are nutrient, metabolic, and hormonal changes, along with physical damage to the uterus occurring during parturition. Maladaptation during the transition period can result in suboptimal immune status, increased disease risks, and impaired reproduction (Ospina et al., 2010; Ster et al., 2012; Wankhade et al., 2017; Trevisi and Minuti, 2018; Pascottini and LeBlanc, 2020). Thus, it is not surprising that nulliparous heifers, with relatively less exposure to the stresses of parturition, lactation, and diseases, have greater conception rates than lactating cows. Peralta et al. (2021) observed that cows showing greater phagocytic capacity in circulating monocytes take longer to conceive. Similarly, Panda et al. (2020) observed that early embryo loss in cows was positively correlated with phagocytic activity, neutrophil/lymphocyte ratio, and expression of proinflammatory cytokines and molecules involved migration in circulating leukocytes. Overall, our findings suggest that cows may have greater neutrophil and other myeloid cell migration, phagocytosis, and proinflammatory signals than heifers. This immunological phenotype may interfere with normal immune adaptations during pregnancy and cause greater embryo loss in cows.

### PBL phenotype of animals that failed to become pregnant

Expression of PPARG, its target gene FABP4, and the anti-inflammatory cytokines IL10 and IL4, were altered in bred-nonpregnant animals compared to cyclic and pregnant cattle. In immune cells, PPARG activation inhibits secretion of proinflammatory cytokines, induces macrophage M2 polarization, promotes neutrophil migration, and ameliorates systemic inflammation observed in metabolic syndrome (Jiang et al., 1998; Fuentes et al., 2013; Abdalla et al., 2020). PPARG agonists enhance the expression of IL10 and IL4 (So et al., 2005; Da Rocha Junior et al., 2013). Moreover, IL4 induces PPARG activation (Paintlia et al., 2006; Jun et al., 2020). Notably, PPARG is involved in regulating insulin sensitivity, glucose and lipid metabolism, and the secretion of cytokines, all of which are important factors for female fertility (Minge et al., 2008). Due to the higher incidence of pregnancy loss in cows with abnormal concentrations of plasma fatty acids, cholesterol, glucose, and indicators of inflammation (Barson et al., 2019; Macmillan et al., 2020; Dirandeh et al., 2021) it is possible that PPARG serves as a regulator of inflammation in PBL during bovine early pregnancy.

Interestingly, the literature suggests a functional antagonism between AHR and PPARG signaling in the biological context of adipogenesis (Alexander et al., 1998), obesity (Shahin et al., 2020) and in the differentiation of T regulatory and Th17 phenotype of lymphocytes (Klotz et al., 2009; Da Rocha Junior et al., 2013; Cipolletta et al., 2015). Analysis of binding sites for transcription factors on promoter regions of the human genome using UCSC Genome Browser (https://genome.ucsc.edu/) and JASPAR database (https://jaspar.genereg.net/) indicate that interferon-induced transcription factors belonging to the interferon regulatory factor (IRF) family and signal transducer and activator of transcription (STAT) family can activate gene expression of both AHR and PPARG. Moreover, PPARG expression can be regulated by the dimers of aryl hydrocarbon receptor nuclear translocator (ARNT) with AHR or with hypoxia inducible factor 1 subunit alpha (HIF1A). On the other hand, the promoter region of AHR lacks binding sites for PPARG. This intricate relationship between PPARG and AHR and their roles in modulating inflammatory responses and various physiological processes require further investigation to understand the role these molecules may play in leukocyte function during pregnancy.

### Conclusions

Our results support the hypothesis that there are differences in the peripheral immune response to pregnancy between cows and heifers. Because cows have lower conception rates than heifers, it is possible that differences in their immune response to pregnancy may play a role in embryo loss. Additionally, cows that fail to become pregnant exhibit immune changes consistent with impaired anti-inflammatory mechanisms in circulating leukocytes. Finally, the balance between pro- and anti-inflammatory response may involve AHR and PPARG signaling, respectively. These findings highlight the role of the immune system in influencing the establishment of pregnancy. What remains to be determined is the cause(s) of immune activation in PBL of cows and if they directly induce pregnancy loss.

## Acknowledgements

This project was supported by USDA Grant no. 2017-67015-26455 and by the NIH-T32 fellowship Grant no. T32GM108563. We thank Dr. Joy Pate for her thoughtful critique of this manuscript and Nadine Houck and Travis Edwards for assistance in preparing and sampling the animals used in these experiments. We thank Ty Montgomery for helping with animal management, sampling, and blood collection in experiment 3.

## Author contributions

TO conceived the research question and experimental design. MIS, NO, FG conducted the experiments. MIS and NO performed data analysis. MIS wrote the manuscript. TO provided significant intellectual contributions during manuscript content review. All authors read and approved the submitted version.

## Supplementary data

Data for the flow cytometry gating strategy, antibody specificity, and the expression of immune molecules from heifers in experiment 1 and cows in experiment 2 showing no significant effects of status (cyclic vs pregnant) and day of sampling can be accessed in the link: https://data.mendeley.com/preview/32cx2t8hhz?a=a1ac84ba-4207-4b3f-a348-f09b1f9ea2c0

## References

Abdalla, H.B., M.H. Napimoga, A.H. Lopes, A.G. de Macedo Maganin, T.M. Cunha, T.E. Van Dyke, and J.T. Clemente Napimoga. 2020. Activation of PPAR-γ induces macrophage polarization and reduces neutrophil migration mediated by heme oxygenase 1. Int Immunopharmacol 8: 106565. 10.1016/j.intimp.2020.106565.

Ahmadi, M.R., S. Nazifi, and H.R. Ghaisari. 2006. Comparison of hormonal changes of estrous cycle with cytology of cervical mucosa and hematological parameters in dairy heifers. Comp Clin Path 15:94–97. 10.1007/s00580-006-0613-7.

Alexander, D.L., L.G. Ganem, P. Fernandez-Salguero, F. Gonzalez, and C.R. Jefcoate. 1998. Aryl-hydrocarbon receptor is an inhibitory regulator of lipid synthesis and of commitment to adipogenesis. J Cell Sci 111:3311–3322. 10.1242/jcs.111.22.3311.

Alhussien, M.N., and A.K. Dang. 2019. Potential roles of neutrophils in maintaining the health and productivity of dairy cows during various physiological and physiopathological conditions: a review. Immunol Res 67:21–38. 10.1007/s12026-019-9064-5.

De Almeida, A.C., S.M. Barbosa, M. De Lourdes Rios Barjas-Castro, S.T. Olalla-Saad, and A. Condino-Neto. 2012. IFN-β, IFN-γ, and TNF-α decrease erythrophagocytosis by human monocytes independent of SIRP-α or SHP-1 expression. Immunopharmacol Immunotoxicol 34:1054–1059. 10.3109/08923973.2012.697470.

Al-ofi, E., S.B. Coffelt, and D.O. Anumba. 2012. Monocyte subpopulations from pre-eclamptic patients are abnormally skewed and exhibit exaggerated responses to toll-like receptor ligands. PLoS One 7: e42217. 10.1371/journal.pone.0042217.

Baban, B., P. Chandler, D. McCool, B. Marshall, D.H. Munn, and A.L. Mellor. 2004. Indoleamine 2,3-dioxygenase expression is restricted to fetal trophoblast giant cells during murine gestation and is maternal genome specific. J Reprod Immunol 61:67–77. 10.1016/j.jri.2003.11.003.

Bahl, K., A. Hüebner, R.J. Davis, and R.M. Welsh. 2010. Analysis of Apoptosis of Memory T Cells and Dendritic Cells during the Early Stages of Viral Infection or Exposure to Toll-Like Receptor Agonists. J Virol 84:4866–4877. 10.1128/jvi.02571-09.

Barbalat, R., L. Lau, R.M. Locksley, and G.M. Barton. 2009. Toll-like receptor 2 on inflammatory monocytes induces type I interferon in response to viral but not bacterial ligands. Nat Immunol 10:1200–1209. doi:10.1038/ni.1792.

Barson, R.K., S. Padder, A.S.M. Sayam, M.M. Rahman, M. Musharraf, U. Bhuiyan, and J. Bhattacharjee. 2019. Serum glucose, urea nitrogen, cholesterol, and total proteins in crossbred repeat breeder and normally cyclic cows. J Adv Vet Anim Res 6:82–85. 10.5455/javar.2019.f316.

Bazer, F.W., T.E. Spencer, and T.L. Ott. 1997. Interferon tau: A novel pregnancy recognition signal. American Journal of Reproductive Immunology 37:412–420. 10.1111/j.1600-0897.1997.tb00253.x.

Belardelli, F. 1995. Role of interferons and other cytokines in the regulation of the immune response. APMIS 103:161–179. 10.1111/j.1699-0463.1995.tb01092.x.

Blumenreich, M.S. 1990. The White Blood Cell and Differential Count. 3rd ed. Butterworths.

Bock, K.W. 2021. Aryl hydrocarbon receptor (AHR), integrating energy metabolism and microbial or obesity-mediated inflammation. Biochem Pharmacol 184:114346. 10.1016/j.bcp.2020.114346.

Botezatu, A., C. Vlagioiu, M. Codreanu, and A. Oraşanu. 201 Biochemical and Hematological Profile in Cattle Effective. Bull Univ Agric Sci Vet Med 71:27–30. 10.15835/buasvmcn-vm:71:1:9544.

Bozzano, F., A. Picciotto, P. Costa, F. Marras, V. Fazio, I. Hirsch, D. Olive, L. Moretta, and A. de Maria. 2011. Activating NK cell receptor expression/function (NKp30, NKp46, DNAM-1) during chronic viraemic HCV infection is associated with the outcome of combined treatment. Eur J Immunol 41:2905–2914. 10.1002/eji.201041361.

Bunaciu, R.P., H.A. Jensen, R.J. MacDonald, D.H. La Tocha, J.D. Varner, and A. Yen. 2015. 6-formylindolo(3,2-b)carbazole(FICZ) modulates the signalsome responsible for RA-induced differentiation of HL-60 myeloblastic leukemia cells. PLoS One 10:e0135668. 10.1371/journal.pone.0135668.

Burnett, T.A., A.M.L. Madureira, B.F. Silper, A. Tahmasbi, A. Nadalin, D.M. Veira, and R.L.A. Cerri. 2015. Relationship of concentrations of cortisol in hair with health, biomarkers in blood, and reproductive status in dairy cows. J Dairy Sci 98:4414–4426. 10.3168/jds.2014-8871.

Burton, J.L., S.A. Madsen, L.C. Chang, P.S.D. Weber, K.R. Buckham, R. Van Dorp, M.C. Hickey, and B. Earley. 2005. Gene expression signatures in neutrophils exposed to glucocorticoids: A new paradigm to help explain “neutrophil dysfunction” in parturient dairy cows. Vet Immunol Immunopathol 105:197–219. 10.1016/j.vetimm.2005.02.012.

Castro, F., A.P. Cardoso, R.M. Gonçalves, K. Serre, and M.J. Oliveira. 2018. Interferon-gamma at the crossroads of tumor immune surveillance or evasion. Front Immunol 9:847. 10.3389/fimmu.2018.00847.

Chawla-Sarkar, M., D.J. Lindner, Y.-F. Liu, B.R. Williams, G.C. Sen, R.H. Silverman, and E.C. Borden. 2003. Apoptosis and interferons: Role of interferon-stimulated genes as mediators of apoptosis. Apoptosis 8:237–249. 10.1023/a:1023668705040.

Cheng, L., H. Yu, G. Li, F. Li, J. Ma, J. Li, L. Chi, L. Zhang, and L. Su. 2017. Type I interferons suppress viral replication but contribute to T cell depletion and dysfunction during chronic HIV-1 infection. J Clin Invest 2:e94366. 10.1172/jci.insight.94366.

Cipolletta, D., P. Cohen, B.M. Spiegelman, C. Benoist, and D. Mathis. 2015. Appearance and disappearance of the mRNA signature characteristic of Treg cells in visceral adipose tissue: Age, diet, and PPARγ effects. Proc Natl Acad Sci U S A 112:482–487. 10.1073/pnas.1423486112.

Dagenais-Lussier, X., H. Loucif, A. Murira, X. Laulhé, S. Stäger, A. Lamarre, and J. van Grevenynghe. 2018. Sustained IFN-I expression during established persistent viral infection: A “bad seed” for protective immunity. Viruses 10:12. 10.3390/v10010012.

Dirandeh, E., M.A. Sayyar, Z. Ansari-Pirsaraei, H. Deldar, and W.W. Thatcher. 2021. Peripheral leucocyte molecular indicators of inflammation and oxidative stress are altered in dairy cows with embryonic loss. Sci Rep 11:12771. 10.1038/s41598-021-91535-2.

Diskin, M.G., S.M. Waters, M.H. Parr, and D.A. Kenny. 2016. Pregnancy losses in cattle: Potential for improvement. Reprod Fertil Dev 28:83–93. 10.1071/RD15366.

Fuentes, E., L. Guzmán-Jofre, R. Moore-Carrasco, and I. Palomo. 2013. Role of PPARs in inflammatory processes associated with metabolic syndrome (Review). Mol Med Rep 8:1611–1616. 10.3892/mmr.2013.1714.

Gargaro, M., G. Manni, G. Scalisi, P. Puccetti, and F. Fallarino. 2021. Tryptophan metabolites at the crossroad of immune-cell interaction via the aryl hydrocarbon receptor: Implications for tumor immunotherapy. Int J Mol Sci 22:4644. 10.3390/ijms22094644.

George, J.W., J. Snipes, and V.M. Lane. 2010. Comparison of bovine hematology reference intervals from 1957 to 2006. Vet Clin Pathol 39:138–148. 10.1111/j.1939-165X.2009.00208.x.

Girer, N.G., C.R. Tomlinson, and C.J. Elferink. 2021. The aryl hydrocarbon receptor in energy balance: The road from dioxin-induced wasting syndrome to combating obesity with AHR ligands. Int J Mol Sci 22:49. 10.3390/ijms22010049.

Green, J.C., C.S. Okamura, S.E. Poock, and M.C. Lucy. 2010. Measurement of interferon-tau (IFN-τ) stimulated gene expression in blood leukocytes for pregnancy diagnosis within 18-20d after insemination in dairy cattle. Anim Reprod Sci 121:24–33. 10.1016/j.anireprosci.2010.05.010.

Groebner, A.E., I. Rubio-Aliaga, K. Schulke, H.D. Reichenbach, H. Daniel, E. Wolf, H.H.D. Meyer, and S.E. Ulbrich. 2011. Increase of essential amino acids in the bovine uterine lumen during preimplantation development. Reproduction 141:685–695. 10.1530/rep-10-0533.

Herman, N., C. Trumel, A. Geffré, J.P. Braun, M. Thibault, F. Schelcher, and N. Bourgès-Abella. 2018. Hematology reference intervals for adult cows in France using the Sysmex XT-2000iV analyzer. J Vet Diagn Invest 30:678–687. 10.1177/1040638718790310.

Huang, T., J. Zhou, and J. Wang. 2022. Calcium and calcium-related proteins in endometrial cancer: opportunities for pharmacological intervention. Int J Biol Sci 18:1065–1078. 10.7150/ijbs.68591.

Huber, J.P., and J. David Farrar. 2011. Regulation of effector and memory T-cell functions by type I interferon. Immunology 132:466–474. 10.1111/j.1365-2567.2011.03412.x.

Hughes, C. H. K., Rogus, A., & Pate, J. L. (2021). REPRODUCTION RESEARCH NR5A2 and potential regulatory miRNAs in the bovine CL during early pregnancy. Reproduction, 161, 173–182. 10.1530/REP

Jaitin, D.A., and G. Schreiber. 2007. Upregulation of a small subset of genes drives type I interferon-induced antiviral memory. J Interferon Cytokine Res 27:653–664. 10.1089/jir.2006.0162.

Jiang, C., A.T. Ting, and B. Seed. 1998. PPAR-gamma agonists inhibit production of monocyte inflammatory cytokines. Nature 391:82–86. 10.1038/34184.

Jun, I., B.R. Kim, S.Y. Park, H. Lee, J. Kim, E.K. Kim, K.Y. Seo, and T. im Kim. 2020. Interleukin-4 stimulates lipogenesis in meibocytes by activating the STAT6/PPARγ signaling pathway. Ocul Surf 18:575–582. 10.1016/j.jtos.2020.04.015.

Kamada, R., W. Yang, Y. Zhang, M.C. Patel, Y. Yang, R. Ouda, A. Dey, Y. Wakabayashi, K. Sakaguchi, T. Fujita, T. Tamura, J. Zhu, and K. Ozato. 2018. Interferon stimulation creates chromatin marks and establishes transcriptional memory. PNAS 115:E9162–E9171. 10.1073/pnas.1720930115.

Kamat, M.M., S. Vasudevan, S.A. Maalouf, D.H. Townson, J.L. Pate, and T.L. Ott. 2016. Changes in Myeloid Lineage Cells in the Uterus and Peripheral Blood of Dairy Heifers During Early Pregnancy. Biol Reprod 95:68,1–12. 10.1095/biolreprod.116.141069.

Klotz, L., S. Burgdorf, I. Dani, K. Saijo, J. Flossdorf, S. Hucke, J. Alferink, N. Novak, M. Beyer, G. Mayer, B. Langhans, T. Klockgether, A. Waisman, G. Eberl, J. Schultze, M. Famulok, W. Kolanus, C. Glass, C. Kurts, and P.A. Knolle. 2009. The nuclear receptor PPARγ selectively inhibits Th17 differentiation in a T cell-intrinsic fashion and suppresses CNS autoimmunity. J Exp Med 206:2079–2089. 10.1084/jem.20082771.

Lee, J.Y., and K.E. Sullivan. 2001. Gamma interferon and lipopolysaccharide interact at the level of transcription to induce tumor necrosis factor alpha expression. Infect Immun 69:2847–2852. 10.1128/IAI.69.5.2847-2852.2001.

Leung, S.T., K. Derecka, G.E. Mann, A.P.F. Flint, and D.C. Wathes. 2000. Uterine lymphocyte distribution and interleukin expression during early pregnancy in cows. J Reprod Fertil 119:25–33. 10.1530/reprod/119.1.25.

Libermann, T.A., and D. Baltimore. 1990. Activation of interleukin-6 gene expression through the NF-kappa B transcription factor. Mol Cell Biol 10:2327–2334, 10.1128/mcb.10.5.2327-2334.1990.

Luppi, P., C. Haluszczak, D. Betters, C.A.H. Richard, M. Trucco, and J.A. Deloia. 2002. Monocytes are progressively activated in the circulation of pregnant women. J Leukoc Biol 72:874–84. 10.1189/jlb.72.5.874.

Macmillan, K., M. Gobikrushanth, I.L. Helguera, A. Behrouzi, and M.G. Colazo. 2020. Relationships between early postpartum nutritional and metabolic profiles and subsequent reproductive performance of lactating dairy cows. Theriogenology 151:52–57. 10.1016/j.theriogenology.2020.03.034.

Mansouri-Attia, N., L.J. Oliveira, N. Forde, A.G. Fahey, J.A. Browne, J.F. Roche, O. Sandra, P. Reinaud, P. Lonergan, and T. Fair. 2012. Pivotal role for monocytes/macrophages and dendritic cells in maternal immune response to the developing embryo in cattle. Biol Reprod 87:123,1–12. 10.1095/biolreprod.112.101121.

McDougall, S., S.J. LeBlanc, and A. Heiser. 2017. Effect of prepartum energy balance on neutrophil function following pegbovigrastim treatment in periparturient cows. J Dairy Sci 100:7478–7492. 10.3168/jds.2017-12786.

McLoughlin, R.M., J. Witowski, R.L. Robson, T.S. Wilkinson, S.M. Hurst, A.S. Williams, J.D. Williams, S. Rose-John, S.A. Jones, and N. Topley. 2003. Interplay between IFN-γ and IL-6 signaling governs neutrophil trafficking and apoptosis during acute inflammation. Journal of Clinical Investigation 112:598–607. 10.1172/JCI17129.

Melgert, B.N., F. Spaans, T. Borghuis, P.A. Klok, B. Groen, A. Bolt, P. de Vos, M.G. van Pampus, T.Y. Wong, H. van Goor, W.W. Bakker, and M.M. Faas. 2012. Pregnancy and Preeclampsia Affect Monocyte Subsets in Humans and Rats. PLoS One 7: e45229. 10.1371/journal.pone.0045229.

Mellor, A.L., and D.H. Munn. 2004. IDO expression by dendritic cells: Tolerance and tryptophan catabolism. Nat Rev Immunol 4:762–77. 10.1038/nri1457.

Mezrich, J.D., J.H. Fechner, X. Zhang, B.P. Johnson, W.J. Burlingham, and C.A. Bradfield. 2010. An Interaction between Kynurenine and the Aryl Hydrocarbon Receptor Can Generate Regulatory T Cells. J Immunol 185:3190–3198. 10.4049/jimmunol.0903670.

Minge, C.E., R.L. Robker, and R.J. Norman. 2008. PPAR gamma: Coordinating metabolic and immune contributions to female fertility. PPAR Res 2008:243791. 10.1155/2008/243791.

Miwa, N., S. Hayakawa, S. Miyazaki, S. Myojo, Y. Sasaki, M. Sakai, O. Takikawa, and S. Saito. 2006. IDO expression on decidual and peripheral blood dendritic cells and monocytes/macrophages after treatment with CTLA-4 or interferon-γ increase in normal pregnancy but decrease in spontaneous abortion. Mol Hum Reprod 11:865–870. 10.1093/molehr/gah246.

Mohapatra, S.K., B.S.K. Panda, A.K. Verma, R. Kapila, and A.K. Dang. 2020. Implantation associated changes in expression profile of indoleamine-2, 3-dioxygenase 1, Th1-Th2 cytokines and interferon-stimulated genes on neutrophils and peripheral blood mononuclear cells of crossbred cows. J Reprod Immunol 142:103188. 10.1016/j.jri.2020.103188.

Moretti, P., M. Probo, A. Cantoni, S. Paltrinieri, and A. Giordano. 2016. Fluctuation of neutrophil counts around parturition in Holstein dairy cows with and without retained placenta. Res Vet Sci 107:207–212. 10.1016/j.rvsc.2016.06.015.

Myers, L.M., M.C. Tal, L.B. Torrez Dulgeroff, A.B. Carmody, R.J. Messer, G. Gulati, Y.Y. Yiu, M.M. Staron, C.L. Angel, R. Sinha, M. Markovic, E.A. Pham, B. Fram, A. Ahmed, A.M. Newman, J.S. Glenn, M.M. Davis, S.M. Kaech, I.L. Weissman, and K.J. Hasenkrug. 2019. A functional subset of CD8 + T cells during chronic exhaustion is defined by SIRPα expression. Nat Commun 10:794. 10.1038/s41467-019-08637-9.

Oliveira, L.J., and P.J. Hansen. 2008. Deviations in populations of peripheral blood mononuclear cells and endometrial macrophages in the cow during pregnancy. Reproduction 136:481–490. 10.1530/REP-08-0218.

Oliveira, L.J., N. Mansourri-Attia, A.G. Fahey, J. Browne, N. Forde, J.F. Roche, P. Lonergan, and T. Fair. 2013. Characterization of the Th Profile of the Bovine Endometrium during the Oestrous Cycle and Early Pregnancy. PLoS One 8:e75571. 10.1371/journal.pone.0075571.

Ospina, P.A., D. V. Nydam, T. Stokol, and T.R. Overton. 2010. Evaluation of nonesterified fatty acids and β-hydroxybutyrate in transition dairy cattle in the northeastern United States: Critical thresholds for prediction of clinical diseases. J Dairy Sci 93:546–554. 10.3168/jds.2009-2277.

Ott, T.L. 2019. Symposium review: Immunological detection of the bovine conceptus during early pregnancy. J Dairy Sci 102:3766–3777. 10.3168/jds.2018-15668.

Paintlia, A.S., M.K. Paintlia, I. Singh, and A.K. Singh. 2006. IL-4-Induced Peroxisome Proliferator-Activated Receptor γ Activation Inhibits NF-κB Trans Activation in Central Nervous System (CNS) Glial Cells and Protects Oligodendrocyte Progenitors under Neuroinflammatory Disease Conditions: Implication for CNS-Demyelinating Diseases. J Immunol 176:4385–4398. 10.4049/jimmunol.176.7.4385.

Panda, B.S.K., S.K. Mohapatra, A.K. Verma, A. Kamboj, M.N. Alhussien, and A.K. Dang. 2020. A comparative study on various immunological parameters influencing embryo survivability in crossbred dairy cows. Theriogenology 157:140–148. 10.1016/j.theriogenology.2020.05.041.

Partida-Sánchez, S., D.A. Cockayne, S. Monard, E.L. Jacobson, N. Oppenheimer, B. Garvy, K. Kusser, S. Goodrich, M. Howard, A. Harmsen, T.D. Randall, and F.E. Lund. 2001. Cyclic ADP-ribose production by CD38 regulates intracellular calcium release, extracellular calcium influx and chemotaxis in neutrophils and is required for bacterial clearance in vivo. Nat Med 7:1209–1216. 10.1038/nm1101-1209.

Pascottini, O.B., and S.J. LeBlanc. 2020. Modulation of immune function in the bovine uterus peripartum. Theriogenology 150:193–200. 10.1016/j.theriogenology.2020.01.042.

Pedersen, H.S., G. Mazzoni, L. Stroebech, H.N. Kadarmideen, P. Hyttel, and H. Callesen. 2017. Basic and practical aspects of pregnancy establishment in cattle. Anim Reprod 14:581–588. 10.21451/1984-3143-AR1001.

Peralta, M.B., S. Cainelli, A.F. Stassi, E. Angeli, M.S. Renna, M.L. Signorini, N.C. Gareis, L. Durante, F. Rey, H.H. Ortega, N.R. Salvetti, and M.M.L. Velázquez. 2021. Association between phagocytic activity of monocytes and days to conception after parturition in dairy cows when considering the hormonal and metabolic milieu. Anim Reprod Sci 232:106818. 10.1016/j.anireprosci.2021.106818.

Peters, M.W., and J.R. Pursley. 2002. Fertility of lactating dairy cows treated with ovsynch after presynchronization injections of PGF2α and GnRH. J Dairy Sci 85:2403–2406. 10.3168/jds.S0022-0302(02)74322-1.

Pohjanvirta, R. 2017. AHR in energy balance regulation. Curr Opin Toxicol 2:8–14 10.1016/j.cotox.2017.01.002.

Da Rocha Junior, L.F., A.T. Dantas, Â.L.B.P. Duarte, M.J.B. De Melo Rego, I.D.R. Pitta, and M.G.D.R. Pitta. 2013. PPAR γ agonists in adaptive immunity: What do immune disorders and their models have to tell us?. PPAR Res 2013:519724. 10.1155/2013/519724.

Rutigliano, H.M., K.A. Leppo, and K.P. Morgado. 2022. Changes in mononuclear immune cells during bovine pregnancy. Reprod Fertil Dev 34:608–618. 10.1071/RD21161.

Le Saout, C., R.B. Hasley, H. Imamichi, L. Tcheung, Z. Hu, M.A. Luckey, J.H. Park, S.K. Durum, M. Smith, A.W. Rupert, M.C. Sneller, H.C. Lane, and M. Catalfamo. 2014. Chronic Exposure to Type-I IFN under Lymphopenic Conditions Alters CD4 T Cell Homeostasis. PLoS Pathog 10: e1003976. 10.1371/journal.ppat.1003976.

Shahin, N.N., G.T. Abd-Elwahab, A.A. Tawfiq, and H.M. Abdelgawad. 2020. Potential role of aryl hydrocarbon receptor signaling in childhood obesity. Biochim Biophys Acta Mol Cell Biol Lipids 1865:158714. 10.1016/j.bbalip.2020.158714.

Shinde, R., K. Hezaveh, M.J. Halaby, A. Kloetgen, A. Chakravarthy, T. Da Silva Medina, R. Deol, K.P. Manion, Y. Baglaenko, M. Eldh, S. Lamorte, D. Wallace, S.B. Chodisetti, B. Ravishankar, H. Liu, K. Chaudhary, D.H. Munn, A. Tsirigos, M. Madaio, S. Gabrielsson, Z. Touma, J. Wither, D.D. De Carvalho, and T.L. McGaha. 2018. Apoptotic cell-induced AhR activity is required for immunological tolerance and suppression of systemic lupus erythematosus in mice and humans article. Nat Immunol 19:571–582. 10.1038/s41590-018-0107-1.

So, R.K., S.L. Kyung, S.P. Hee, J.P. Seoung, H.M. Kyung, M.J. Sun, and C.L. Yong. 2005. Involvement of IL-10 in peroxisome proliferator-activated receptor γ-mediated anti-inflammatory response in asthma. Mol Pharmacol 68:1568–1575. 10.1124/mol.105.017160.

Sojka, D.K., L. Yang, B. Plougastel-Douglas, D.A. Higuchi, B.A. Croy, and W.M. Yokoyama. 2018. Cutting Edge: Local Proliferation of Uterine Tissue-Resident NK Cells during Decidualization in Mice. J Immunol 201:2551–2556. 10.4049/jimmunol.1800651.

Solano, M.E. 2019. Decidual immune cells: Guardians of human pregnancies. Best Pract Res Clin Obstet Gynaecol 60:3–16. 10.1016/j.bpobgyn.2019.05.009.

Spencer, T.E. 2013. Early pregnancy: Concepts, challenges, and potential solutions. Anim Front 3:48–55. 10.2527/af.2013-0033.

Ster, C., M.C. Loiselle, and P. Lacasse. 2012. Effect of postcalving serum nonesterified fatty acids concentration on the functionality of bovine immune cells. J Dairy Sci 95:708–717. 10.3168/jds.2011-4695.

Stobbe-Maicherski, N., S. Wolff, C. Wolff, J. Abel, U. Sydlik, K. Frauenstein, and T. Haarmann-Stemmann. 2013. The interleukin-6-type cytokine oncostatin M induces aryl hydrocarbon receptor expression in a STAT3-dependent manner in human HepG2 hepatoma cells. FEBS Journal 280:6681–6690. 10.1111/febs.12571.

Trevisi, E., and A. Minuti. 2018. Assessment of the innate immune response in the periparturient cow. Res Vet Sci 116:47–54. 10.1016/j.rvsc.2017.12.001.

Vasudevan, S., M.M. Kamat, S.S. Walusimbi, J.L. Pate, and T.L. Ott. 2017. Effects of early pregnancy on uterine lymphocytes and endometrial expression of immune-regulatory molecules in dairy heifers. Biol Reprod 97:104–118. 10.1093/biolre/iox061.

Wankhade, P.R., A. Manimaran, A. Kumaresan, S. Jeyakumar, K.P. Ramesha, V. Sejian, D. Rajendran, and M.R. Varghese. 2017. Metabolic and immunological changes in transition dairy cows: A review. Vet World 10:1367–1377. 10.14202/vetworld.2017.1367-1377.

Welsh, R.M., K. Bahl, H.D. Marshall, and S.L. Urban. 2012. Type 1 interferons and antiviral CD8 T-Cell responses. PLoS Pathog 8: e1002352. 10.1371/journal.ppat.1002352.

Whitmire, J.K., J.T. Tan, and J.L. Whitton. 2005. Interferon-γ acts directly on CD8+ T cells to increase their abundance during virus infection. J Exp Med 201:1053–1059. 10.1084/jem.20041463.

Yang, L., Y. Wang, X. Ma, S. Wang, and L. Zhang. 2016. Changes in expression of Th1 and Th2 cytokines in bovine peripheral blood mononuclear cells during early pregnancy. Indian J Anim Res 50:466–470. 10.18805/ijar.5538.

Ye, Z.H., X.M. Jiang, M.Y. Huang, Y.L. Xu, Y.C. Chen, L.W. Yuan, C.Y. Huang, W.B. Yu, X. Chen, and J.J. Lu. 2021. Regulation of CD47 expression by interferon-gamma in cancer cells. Transl Oncol 14:101162. 10.1016/j.tranon.2021.101162.

Zhang, X., S. SUn, I. Hwang, D.F. Tough, and J. Sprent. 1998. Potent and Selective Stimulation of Memory-Phenotype CD8 T Cells In Vivo by IL-15. Immunity 8:591–599. 10.1016/s1074-7613(00)80564-6.

